# On identifying collective displacements in apo-proteins that reveal eventual binding pathways

**DOI:** 10.1101/342253

**Authors:** Dube Dheeraj Prakashchand, Navjeet Ahalawat, Himanshu Khandelia, Jagannath Mondal, Surajit Sengupta

**Affiliations:** Tata Institute of Fundamental Research, Center for Interdisciplinary sciences*, Hyderabad 500107,* India; MEMPHYS, Center for Biomembrane Physics, Department of Physics, Chemistry and Pharmacy, University of Southern Denmark, 5230 Odense M, Denmark

## Abstract

Binding of small molecules to proteins often involves large conformational changes in the latter, which open up pathways to the binding site. Observing and pinpointing these rare events in large scale, all-atom, computations of specific protein-ligand complexes, is expensive and to a great extent serendipitous. Further, relevant collective variables which characterise specific binding or un-binding scenarios are still difficult to identify despite the large body of work on the subject. Here, we show that possible primary and secondary binding pathways can be discovered from short simulations of the apo-protein without waiting for an actual binding event to occur. We use a projection formalism, introduced earlier to study deformation in solids, to analyse local atomic displacements into two mutually orthogonal subspaces — those which are “affine” i.e. expressible as a homogeneous deformation of the native structure, and those which are not. The susceptibility to non-affine displacements among the various residues in the apo-protein is then shown to correlate with typical binding pathways and sites crucial for allosteric modifications. We validate our observation with all-atom computations of three proteins, T4-Lysozyme, Src kinase and Cytochrome P450.

## Introduction

Modern drug discovery operates around the principle of molecular recognition in the context of protein-ligand binding. In the preliminary stages, drug discovery programs often invest in quantifying the binding affinity of suitable drug candidates towards a well-established protein target. However, such efforts often encounter scenarios where identification of the ligand-binding cavity on the protein target becomes challenging. In addition, screening across a large pool of protein receptor candidates as potential drug targets for a specific drug proves tedious and time consuming.

A popular and inexpensive approach for zeroing on a suitable combination of protein receptor and ligand has been drug design based on *in silico* techniques. Towards this end, macromolecular docking^3–7^ based techniques are routinely employed for screening protein-ligand recognition partners. One of the major drawbacks of docking based approach has been the heuristic choice of the scoring function implemented in the docking programs to quantify and rank-order the protein-ligand affinity. Further, the other major criticism has been that in most of the cases the receptor cavity’s inherent flexibility is not taken into account for docking purposes. Rather, docking softwares^8–19^ consider the protein cavity to be rigid and perform the search for a ligand’s ability to dock only in a limited area of protein, often pre-determined by the user’s intuition-based inputs. As a result, docking based drug discovery protocols are fraught often with failures in identifying the correct set of protein ligand combinations.

Molecular Dynamics simulations, with the recent emergence of Graphics-Processing-Unit (GPU) based computation and special purpose machine like Anton, has slowly emerged as an accurate approach to simulate the complete process of protein ligand binding in its atomistic details. The approach, majorly pioneered by Shaw and coworkers^20–23^ has been able to decipher the kinetic pathways leading to spontaneous recognition of the ligand by the receptor. This process is frequently being supplemented by judicious application of Markov State Model^24–26^ based techniques to infer long time recognition processes by shorter time simulations. These techniques being unbiased and devoid of user intervention, currently remain one of the best approaches to explore the protein ligand recognition at their highest resolution. But the process comes at the expense of high computational cost and extended wall clock time. As a result, a plan for practical usage of the Molecular dynamics simulation for the purpose of serious drug discovery is still in its infancy.

The current work offers a practical middle ground between expensive Molecular Dynamics simulation of protein-ligand recognition processes and inaccurate docking based approach by employing an idea from materials physics, which correctly accounts for deformations in protein structure relevant for binding. We use a formalism, introduced earlier to analyse atomic displacements in solids,^27–29^ to predict the protein locations which are potentially susceptible towards ligand recognition. We build on our work under the basic premises that all external and internal stresses result in two types of mutually orthogonal deformations namely those which are “affine” i.e. expressible as a homogeneous deformation of the native structure, and those which are not i.e. ‘non-affine’. As further elaborated in a later section, the current study characterizes the local arrangements of atoms inside protein by probing an important metric termed as “Non Affine parameter” (NAP)^27^. The approach is first convincingly validated by reproducing key conformational changes en route to ligand entry to a mutant T4 Lysozyme. This formalism is then used to propose potential regions which are implicated in “gate-opening” and capture the conformational dynamics during the binding process in the protein receptor Src kinase and Cytochrome P450. Finally, we show how spatial correlations of NAP can elucidate sites for allosteric control in these proteins.

## Protein models and simulation details

We explore two independent protein-ligand systems. Most of the validations of our approach has been done on binding of benzene to the solvent-inaccessible cavity of the L99A mutant of T4 Lysozyme^33^ (T4Lys), a system which has served long as the prototypical protein ligand system for biophysical studies. Some of us had previously performed multi-microsecond molecular dynamics simulation where we have simulated the complete kinetic process of ligand recognition^33^. The details of atomistic simulation process and models for this system have been discussed therein. Apart from the binding process of simulation, in the current work, we have also simulated the apo or ligand-unbound form of T4Lys (PDB id: 3DMV) and and Src kinase (pdb id: 1Y57). Some data for Cytochrome P450 (pdb id: 2CPP) has also been shown. All the apo forms have been modeled using same simulation protocols. We have used the CHARMM36 force field^34^. The protein is solvated by 11613 TIP3P water molecules in a cubical box of dimension 7.18 nm. Sufficient number of ions were added to maintain 150 mM salt concentration. All MD simulations were performed with the Gromacs 5.0.6 simulation package^35^. During the simulation, the average temperature was maintained at 303K using the Nose-Hoover thermostat^36,37^ with a relaxation time of 1.0 ps and an average isotropic pressure of 1 bar was maintained with the Parrinello-Rahman barostat^38^. The Verlet cutoff scheme was employed throughout the simulation with the truncated and shifted Lenard Jones interaction extending to 1.2 nm and long-range electrostatic interactions^39^ treated by Particle Mesh Ewald (PME) summation^40^. All bond lengths involving hydrogen atoms of the protein and the ligand benzene were constrained using the LINCS algorithm^41^ and water hydrogen bonds were fixed using the SETTLE^42^ approach. Simulations were performed using the leapfrog integrator with a time step of 2 fs and initiated by randomly assigning the velocities of all particles. The total time for our simulations of the apo-proteins amounted to (T4Lys=493.54ns, Src kinase=460.00ns, P450=500ns) 1453.54 ns which was sufficient to obtain equilibrated data. It is quite remarkable that our projection formalism yields useful data from such short runs.

## Non Affine Parameter is a novel collective coordinate for protein structure

Atoms of an isolated protein molecule in aqueous solution and at non-zero temperatures undergo continuous random motion. For most proteins, these atomic motions may be analysed in terms of displacement fluctuations about some fixed native structure. Some of these fluctuations represent local affine distortions of the native state while others correspond to non-trivial conformation changes. It is the latter which are associated with functional aspects of the protein. Determining functionally relevant conformational changes is a challenge, especially if such distortions are subtle and have to be extracted from the random background noise. We show now how a collective variable may be defined, which does exactly that. This collective variable, which we call the non-affine parameter, was first defined for crystalline solids. In solids, non-affine displacements are obtained by systematically projecting out^27^ trivial homogeneous deformations of atoms, capturing only those displacements responsible for irreversible plastic events.^28,29^ The NAP, is the squared sum of the mean amplitudes of these displacements. In proteins NAP behaves as a collective coordinate responsible for revealing binding pathways and also for determining possible allosteric couplings between spatially distant residues.

Consider for example (see Fig 1a) an atom *i* and its neighbourhood Ω*_i_* which contains *j* = 1, *n*_Ω_ neighbours. The size of Ω*_i_* is fixed. Too small a Ω*_i_* results in large fluctuations while making Ω*_i_* too large may average out the relevant signal and make it disappear. Typically, we use a neighbourhood which contains 50 – 100 atoms excluding H-atoms. The instantaneous positions of the atoms **r***_i_* at any particular time step are compared with a set of fixed reference positions **R***_i_* taken, for example, from the (ligand-free) native state of the protein. NAP is then defined as the value of the quantity 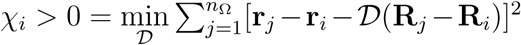 where the minimisation is over choices of the deformation matrix ***D***. NAP is therefore the least square error incurred in trying to represent an arbitrary displacement as an affine or homogeneous deformation of the reference configuration.

**Figure 1:**
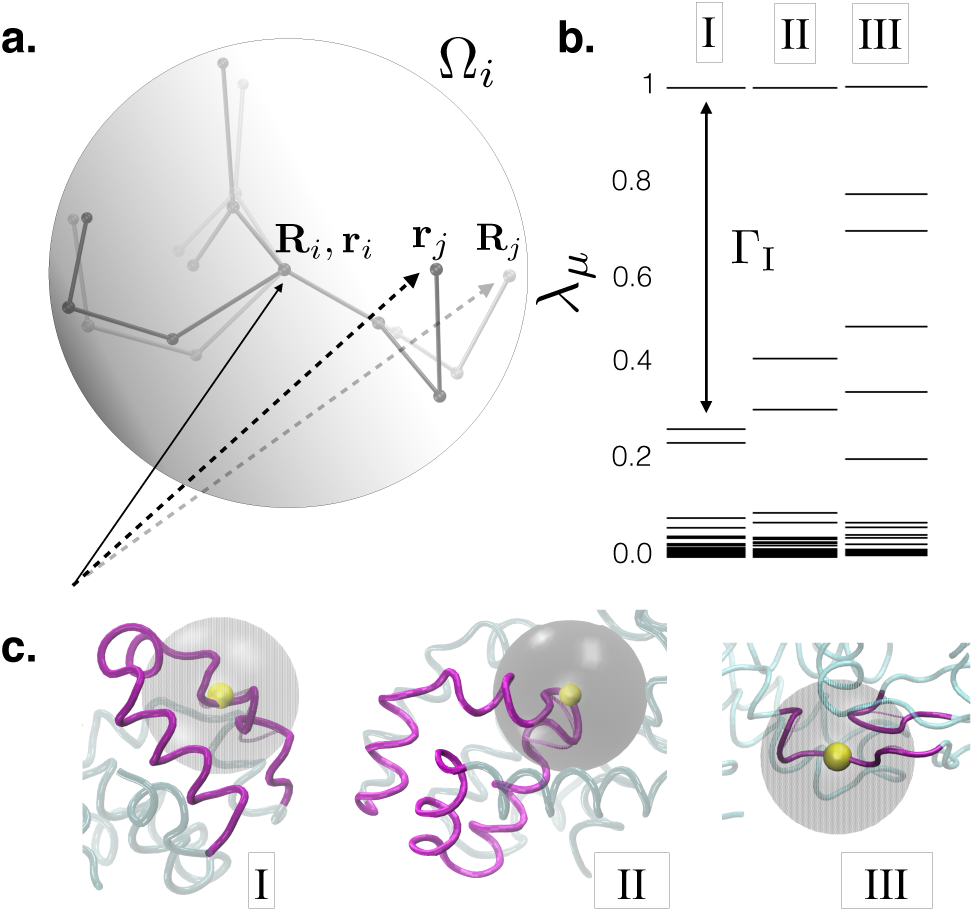
**a.** Schematic diagram used for defining the NAP, *χ_i_* for particle *i* (see text). The neighbourhood Ω*_i_* centred around *i* is shown as a sphere. The reference positions of the neighbours **R***_j_* are shown in light grey while the instantaneous positions **r***_j_* are shown in dark grey. We have subtracted off the displacement of particle *i* (**R***_i_* = r*_i_*) for simplicity. **b.** The distribution of the eigenvalues *λ_μ_* of PCP for three locations in the protein T4Lys, location A corresponds to a carboxyl C atom in helix 7, B corresponds to a carboxyl O atom in helix 5 and C corresponds to an *α* – *C* in a relatively inflexible portion of a *β* sheet. In each of the locations A and B, which are active during ligand binding, the largest eigenvalue is separated from the rest by a large gap Γ*_i_.* The gap is markedly smaller in location C which is relatively inert. **c.** Local structure of the T4Lys protein showing the neighbourhood analysed in **b.** The helices and *β –* sheets are in purple while the rest of the protein is in green. The dominant eigenvector is also shown in transparent purple (see also SI movies V1 & V2.

The minimisation process is actually equivalent^27^ to a projection. The instantaneous displacement of atom *i* is **u***_i_* = **r***_i_* − **R***_i_*. Within the region Ω*_i_,* displacements of neighbouring particles relative to that of *i* are given by **Δ***_ij_* = **u***_j_* − **u***_i_*. Next we define a projection of **Δ***_ij_* onto the non-affine subspace using a projection operator P where P = I − R(R^T^R)^−1^R^T^, and the *n*_Ω_*d* × *d*^2^ elements of **R***_j_*_α,γγ′_ = δ_αγ_*R_j_*_γ′_, taking *i* to be at the origin. Finally, one may show *χ*(**R***_i_*) = **Δ**^T^P**Δ** where we have used the definition **Δ,** as an *n*_Ω_*d* dimensional column vector constructed by rearranging **Δ***_ij_*. This projection formalism can be carried out for any given set of {Ri}. This identification also allows us to compute statistical averages, probability distributions and spatio-temporal correlation functions using standard methods of statistical mechanics. For example, one derives^27^ the equilibrium ensemble average 〈*χ*〉 = ∑*_μ_ λ_μ_* where *λ_μ_* are the eigenvalues of the matrix PCP with **C** = 〈**Δ**^T^**Δ**〉 the local displacement correlator, and we have suppressed the particle index *i* for simplicity. Associated with NAP, we can also define the local NAP susceptibility 〈(*χ_i_* − 〈*χ_i_*〉)^2^〉 and the spatial correlation function 〈(*χ_i_* − 〈*χ_i_*〉)(*χ_j_* − 〈*χ_j_*〉)〉, in the same way as it is done for a crystalline solid.^28^

One of our important results concerns the distribution of the eigenvalues *λ_μ_.* For crystalline solids we had observed^28^ that (1) this distribution is non uniform and the largest eigenvalue is separated from the rest by a large gap and (2) the eigenvector corresponding to the largest eigenvalue represented the dominant plastic mode eg. the creation of a defect dipole. Quite interestingly, the situation is similar here.

In Fig.1b we have plotted the relative magnitudes of the eigenvalues of the local PCP matrix for three different locations on the protein T4Lys. The first one (I) corresponds to an Carbon atom on a Carboxyl group on residue 132 (amino acid Asn) in a region of helix 7, the second one (II) on an Oxygen atom of a Carboxyl group on helix 5 corresponding to residue 105 (amino acid Gln), and the third one (III) for an *α–* Carbon atom within *β* sheet 1, belonging to residue 13, (amino acid Leu). We shall see later that regions I and II are both quite active during binding events while region III is relatively inert. The eigenvalue spectrum surprisingly bears an imprint of this behaviour. The active regions appear to be dominated by a single non-affine mode which is particularly soft. The eigenvalue for this mode is larger than the rest by a substantial amount producing a gape Γ*_i_* in the non-affine spectrum. The inert regions of the proteins have more uniformly distributed eigenvalues. By looking at the structure alone, (see Fig. 1c) it would have been impossible to guess this fact.

In Fig. 2**a** & **b** we show that the distribution of the eigenvalues of the PCP matrix are not random. The soft modes that prove out to be useful in our study correspond to a set of eigenvalues which are quite well separated from the spectrum of the rest of the eigenvalues. In order to demonstrate this, in Fig. 2 **a**, we plot the distribution of the various eigenvalues of the PCP matrix for region (I) of T4Lys (Fig. 1**c**), which is involved in the gate opening resulting from the movements of helices 7 and 9. This is fitted to the Marchenko Pastur (MP) distribution from the prediction from Random Matrix Theory^43–45^. It is clear that the fit is not perfect and there are several eigenvalues which are separated from the rest by gaps. On the other hand, if the matrix PCP was was constructed out of **Δ** which were completely random then we should have obtained the distribution enclosed under the red curve representing the MP distribution for our data. This is shown explicitly for a random correlation matrix in Fig. 2 **b**. The fact that we have a sparsely distributed tail of few relatively very large eigenvalues widely separated from the rest of the spectrum and these large eigenmodes contribute most towards the major conformational changes, clearly demonstrates that our analysis is not a straightforward strain-analysis.^46^ Instead, indeed, there is an underlying physics that can be revealed by focusing on the non-affine subspace P**Δ** of the total displacement **Δ.** Non-affinity may play an important role in guiding a majority of the routine functions performed by the proteins.

**Figure 2:**
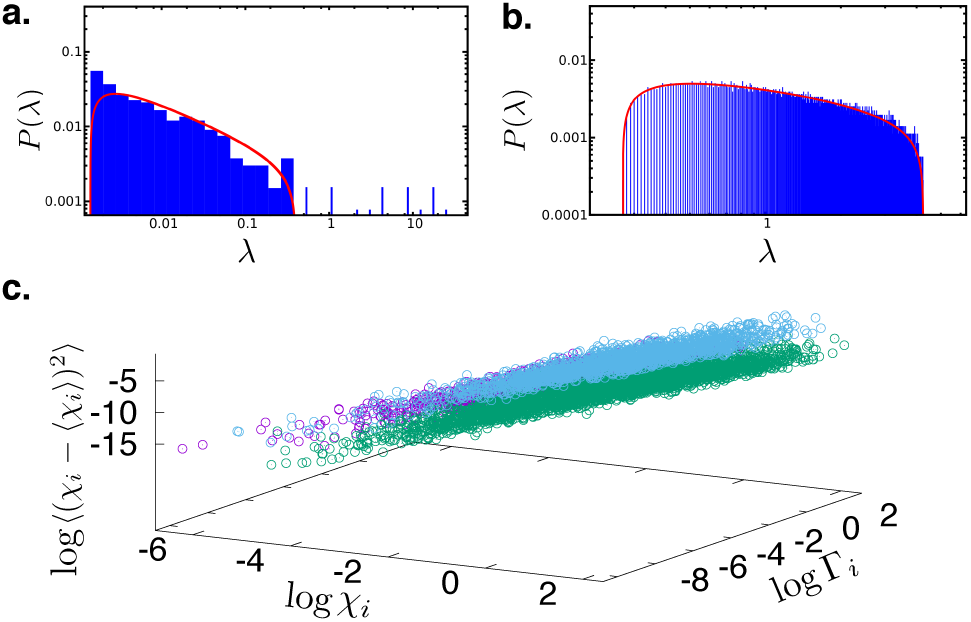
**a.** Distribution of the eigenvalues of the PCP matrix at a specific location in the protein T4Lys (blue histogram) compared with the fitted Marchenko-Pastur distribution (red curve). **b.** A similar fit for a truly random correlation matrix constructed out of vectors with the same number of components as in the first case but now taken from a Gaussian random distribution of zero mean and unit standard deviation.c. Three dimensional plot of the local NAP, *χ_i_,* the gap Γi and the NAP susceptibility for all locations analysed for three different proteins of widely diverse structure and function viz. T4Lys (purple symbols), Src kinase (cyan symbols) and P450 (green symbols). The relation between all three variables for the three proteins is same within the scatter of the data. All the neighbourhoods analysed contained 50 – 100 atoms.

The relation between activity i.e. a large NAP susceptibility and gaps, Γ*_i_*, appears to be an universal feature of all the three proteins, with very diverse structures and functions, studied by us. In Fig, 2**c** we have plotted the NAP susceptibility, Γ*_i_* and NAP for several regions within three proteins T4Lys, P450 and Src kinase. In all of these proteins the three quantities are positively correlated with each other and the nature of the correlation (the slope of the curves) appears to be the same within the error bars for all the proteins.

We demonstrate next that regions of the protein which are involved in a binding event and show large NAP values *during the process of ligand binding* also have large NAP susceptibility in the *apo form of the proteins without the ligand.* The motions of the protein during binding events are also captured by the dominant non-affine eigenmode. Finally we show that large NAP correlations between spatially distant parts of the proteins point out domains of possible allosteric control.

## Non Affine Parameter successfully tracks the ligand binding event in T4 Lysozyme

We first validate the ability of NAP to successfully trace the ligand binding pathway for a recently studied protein-ligand system by Mondal *et al.*^33^, namely benzene binding to T4Lys. Briefly, Mondal *et al.* has recently captured the full details of the benzene binding pathways to the buried cavity of T4Lys using multi-microsecond long atomistic MD simulations. The resulting binding trajectories had identified more than one binding pathways. A key observation from those simulated trajectories was that the benzene binding to the solvent-inaccessible cavity of T4Lys did not require large-scale protein conformational change. Rather, the simulations discovered that it is the subtle displacement of helices of the protein which opens up productive pathways leading to ligand binding in the buried cavity.

Specifically, as plotted in the right hand axis of Fig. 3**a**, a subset of the trajectories pinpointed that prior to benzene binding, the distance between helix-4 and helix-6 increases transiently by around 2 – 3 Å from its equilibrium value, facilitating benzene-gating in the cavity, before it reverts back to its original equilibrium distance after the binding event is complete. On the other hand, another subset of trajectories, spawned from same starting configuration, revealed an alternate pathway of ligand binding, where the transient increase in the distance between a different pair of helices namely, helix-7 and helix-9 triggers the the ligand entry to the cavity, as depicted in Fig. 3**b** (right hand axis). The snapshots shown at the bottom panel of the Fig. 3**a** and **b** captures the location of the benzene relative to the helix gateway during the period of binding event. The three different snapshots labelled as **1, 2** and **3** respectively represent the the helix conformations before, during and after the ligand binding event. The creation of a helix gateway prior to ligand binding (snapshot **2** in each case) is very evident from the increased inter-helix distance.

**Figure 3:**
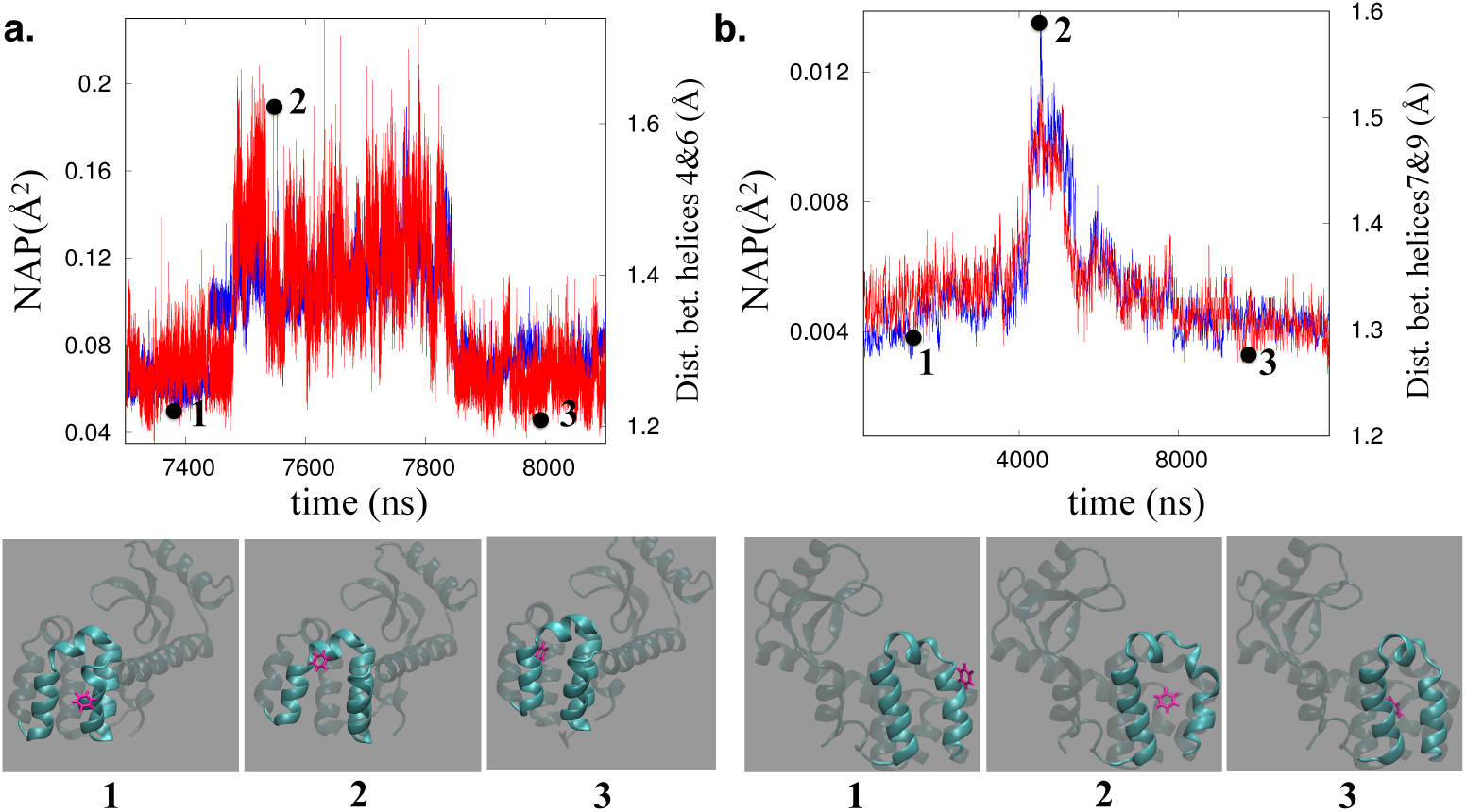
NAP as a good descriptor of ligand binding event: Correlation between time profile of inter-helix distance (blue curves right-hand axis) and corresponding NAP of the helices (red curve, left hand axis) during the process of ligand (benzene) recognition to the buried cavity of L99A T4 Lysozyme. **a.** The trajectory involving helix4-helix6 gateway leading to ligand binding, **b.** the trajectory involving helix7-helix9 gateway leading to ligand recognition. Below each of these curves we also show the ligand (pink) and the protein (green) at three time slices with positions corresponding to (1) outside the gate, (2) within the gate and (3) after the binding event. The three time time slices are also labelled (black dots) in the NAP vs time plots **a.** and **b.**

We calculate the time profile of NAP, averaged over all heavy atoms involving these helices and compare its correspondence with corresponding inter-helix distance. As presented in Fig. 3**a**, we plot the profile of average NAP involving helix-4 and helix-6 on the left hand axis and that of average distance between these two helices on the right hand axis, for the part of time period spanning the benzene entry to the cavity. We find that, both NAP profile and distance profile involving helix-4 and helix-6 follows almost identical trend during the course of ligand entry. Similarly, as shown in Fig. 3**b**, for the other trajectory where ligand binding occurred via helix-7 and helix-9 gateway, time profile of NAP involving helix-7 and helix-9 reproduces the trend of distance fluctuation involving helix-7 and helix-9 quite accurately. The remarkable consistency between the NAP profile and displacement profile of key-profiles responsible for ligand binding suggests that these subtle helix fluctuations are results of non-affine displacements of local environment for each particle. Overall, these results validate the potential of NAP as a suitable collective variable or descriptor for tracking ligand binding kinetics.

## NAP susceptibility of apo protein highlights potential ligand hotspots

The results discussed in previous section have validated the role of NAP as a good descriptor of a ligand binding event and its ability to track the ligand binding pathways. But the validation needed the extensive simulation of actual ligand binding events. As an important finding of the current work, in this section we show that by analysing the NAP susceptibility on a ligand-free (apo) protein itself, one can predict potential protein sites for facile ligand approach, otherwise known as “ligand hotspots”. Since MD simulations of the apo protein do not need to track rare binding events, these cost a small fraction of the time needed for simulating the full protein-ligand systems. Figs. 4 **a** and **c** illustrate the spatial distribution of computed NAP susceptibility of two protein systems in their ligand-free (apo) form, namely, T4Lys and Src kinase. For the purpose of comparison, we have also plotted the locations on protein surface which ligand frequently visits in previously carried out multi-microsecond long unbiased ligand-binding simulations (Figs. 4 **b** and **d**)

**Figure 4:**
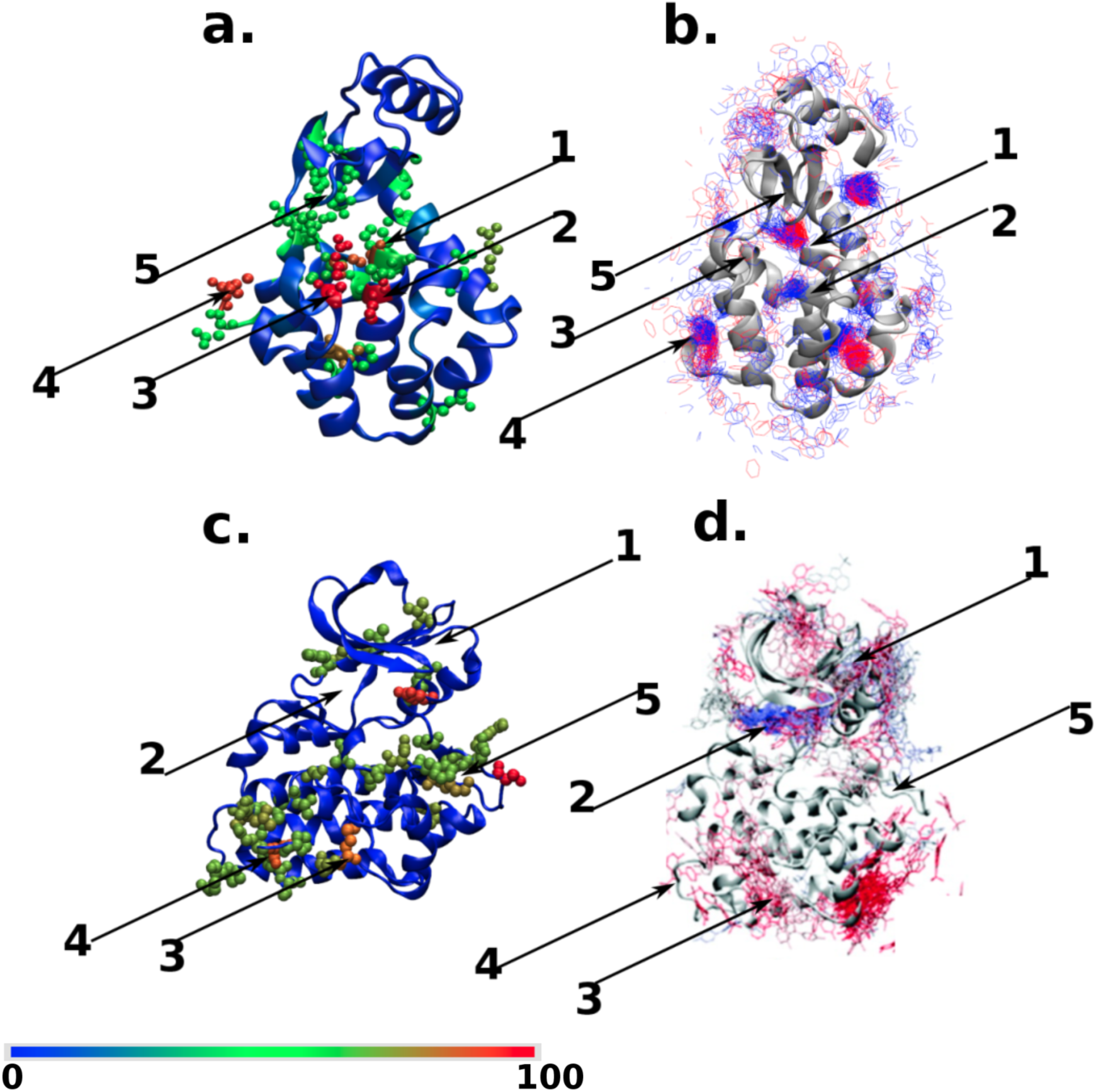
NAP susceptibility of apo proteins (a,c) and ligand binding hotspots (**b**,**d**). **a** The spatial distribution of the NAP susceptibility shown as a colour map red (scaled to 100) to blue (0) for L99A T4 Lysozyme. **b.** In the adjacent panel, we show the same protein now together with snapshots of the ligand. Ligand snapshots in red represents configurations obtained from 4 - 5 *μ*s and blue from 5 - 6 *μ*s of a long MD trajectory. After 6 *μ*s the ligand stays close to the binding gateway. Panels **c** and **d** show the corresponding plots for Src kinase. In **c** the colours have meanings same as in **a**. Panel **d** is reproduced with permission from Ref.^20^, red represents configurations earlier than 15 *μ*s while blue from 15-20 *μ*s. Locations within the protein which are important for ligand binding are numbered and are described in detail in the text. Note that in both cases the regions which have large NAP susceptibility are also those where ligands spend most of their time during binding events.

Fig. 4**a** represents the three-dimensional map of NAP susceptibility of T4Lys. Location 2 of T4Lys includes residues with very high NAP susceptibility (shown in red spheres). A comparison with corresponding location **2** of the time-trace of the ligand in Fig. 4 **b**, as obtained from unbiased ligand-binding trajectories, identifies this location as the native ligand binding site at the buried cavity near the 102*^nd^* residue (a part of helix 5). Further, apart from the final ligand binding location at the 102*^nd^* residue, the computed NAP susceptibilities *around* ligand-free T4Lys were also found to be quite high near locations **1** and **3,** which are respectively the 105*^th^* (a part of helix 5) and 137*^th^* residues (gateway between helix-7 and helix-9). The residues 102 and 105 (which are parts of helix-5) act as hinge-centers or pivots about which the neighboring residues undergo relative re-arrangement opening up the gateway between helices 4 and 6. Similarly, the residue 137 (which is a part of helix-8) acts as a pivot for the gateway between helix 7 and 9. The residue 164 corresponds to location **4** in T4Lys which also has a very high NAP-susceptibility value as it is a terminal residue-part of random coil, mildly contributing towards re-arrangement in helix-9. Interestingly, as shown in Fig. 4**b**, previously simulated independent binding trajectories have also revealed that the pathways leading to ligand entry to the buried cavity of T4Lys involve the helices near locations **1, 2** and **3.** More over, the NAP susceptibility analysis recognises multiple locations with moderate NAP susceptibility (shown by green sphered residues), which are also regularly visited by ligands. Although away from major binding sites, these sites serve as possible precursors to intermediates leading to final molecular recognition. Taken together, the ability to correctly predict eventual ligand binding site and other accessory potential ligand hotspots at T4Lys in its apo form by measuring NAP susceptibility, without the need for prior of knowledge of protein-ligand binding interactions, rates NAP susceptibility as a very promising metric.

The display of the hinge action which opens the gateway to the hydrophobic binding cavity is demonstrated by the supporting movie V1 (opening-up of helix-7 and helix-9) and V2 (opening-up of helix-4 and helix-6), see (*SI Text* for further details). Note that these are not MD simulation snapshots but represent the dominant eigen-displacements arising from a single non-affine mode with the largest eigenvalue. The collective motion involves a “twisting” of the helices and is a very special linear combination of hundreds of local normal modes from which all affine (strain-like) distortions have been systematically projected out. We return to this issue later in the paper.

The ability of NAP susceptibility to predict the potential ligand hotspots on a protein surface is also validated in another popular receptor protein, namely the Src kinase. Fig. 4 **c** shows the NAP susceptibility map around Src kinase in its ligand-free form while the corresponding simulated ligand binding trajectories, as previously simulated by Shaw and coworkers, is shown in Fig. 4 **d**. In this case as well, the ATP-binding site (location **2)** is correctly predicted to have one of the highest NAP susceptibilities in the apo-form and independent simulated ligand-binding trajectories also confirm this. Moreover, other locations of kinase rated as high to intermediate NAP susceptibility for ligand approach are also independently found to be locations regularly traced by ligand in the unbiased ligand binding trajectories by Shaw and coworkers^20^. Specifically, so-called PIF site, MYR site and G-loop site of kinase (PIF-site:region1, MYR-site:region4,), which were found to attract regular ligand visit in actual ligand binding trajectories, are also predicted to have high to moderate NAP susceptibility for the ligand in apo form of the kinase itself. However, interestingly, our NAP susceptibility analysis also predicts a new location **5** deemed highly susceptible to ligand, which is otherwise not known as a ligand hotspot. The discovery of this new site prompted further analysis to justify the observed high susceptibility for the ligand. In the next section, we show that the newly obtained regions of Src kinase which have high NAP-susceptibility also play an important role in establishing an allosteric communication across the protein structure.

## NAP correlations and allostery

In the previous section, we have explored the NAP susceptibilities for different locations of the protein and using this as a metric we were able to extract the important ligand hotspots, in accordance with the simulated ligand binding trajectories. Similarly spatial NAP correlation function or covariance, calculated at the same time, as defined previously, measures the correlation between non-affine displacement fluctuations at two different, possibly distant, locations. We link this quantity to a measure of allostery in the protein systems i.e. the ability of spatially distal sites in proteins to influence catalytic or binding activity..

In Fig. 5, we plot residue-residue NAP covariance matrix representing spatial correlation functions for our two protein systems, namely T4Lys (Fig. 5**a**) and Src kinase (Fig. 5**b**) respectively. On the two axes we have the residue ids and the value of the spatial correlation function between two residues is given by the colour of the point. High values above a suitable cut-off are in yellow and the rest are in black. We are interested in those off-diagonal residues that are spatially far away from each other, since spatially adjacent residues will be trivially correlated due to direct interactions. As shown in Fig. 5**a**, we could get five prominent locations in the matrix for T4Lys which have high NAP correlation at a spatially far-off distance (see *SI Text* for further details of these regions). For example, we found that a very high NAP correlation exists between residue 137 in region **3** and residue 18 in region **5** (labelled (**3,5**)) of T4Lys. This allosteric connection can be seen to be contributing towards regulation of the hydrogen bond between the residue 22 Glu and residue 137 Arg. It is confirmed by Greener et al.^47^ that, using a newly introduced method ExProSE, the breaking of a hydrogen bond between Arg137 and Glu22 causes the opening motion taking the protein from a closed active site cleft conformation to an open active site cleft conformation. Also, earlier, Mchaourab et al.^48^ have confirmed that in solution there is a conformeric equilibrium between substrate-bound and substrate-unbound T4 Lysozyme which is in accord with the active site cleft opening upon substrate removal due to Hinge-Bending motion. Mchaourab et al. used the Site-Directed Spin Labeling to detect these motions. From our NAP formalism, too,^28^ we can see that the softest non-affine modes corresponding to the residues 18 and 137 act as the nucleating-precursors to the opening of the active site cleft. These modes contribute by initially breaking the hydrogen bond between residues 22 and 137 and therefore we can directly see the increasing distance between residue 22 and 137 in video V3 in the supplementary info.

**Figure 5:**
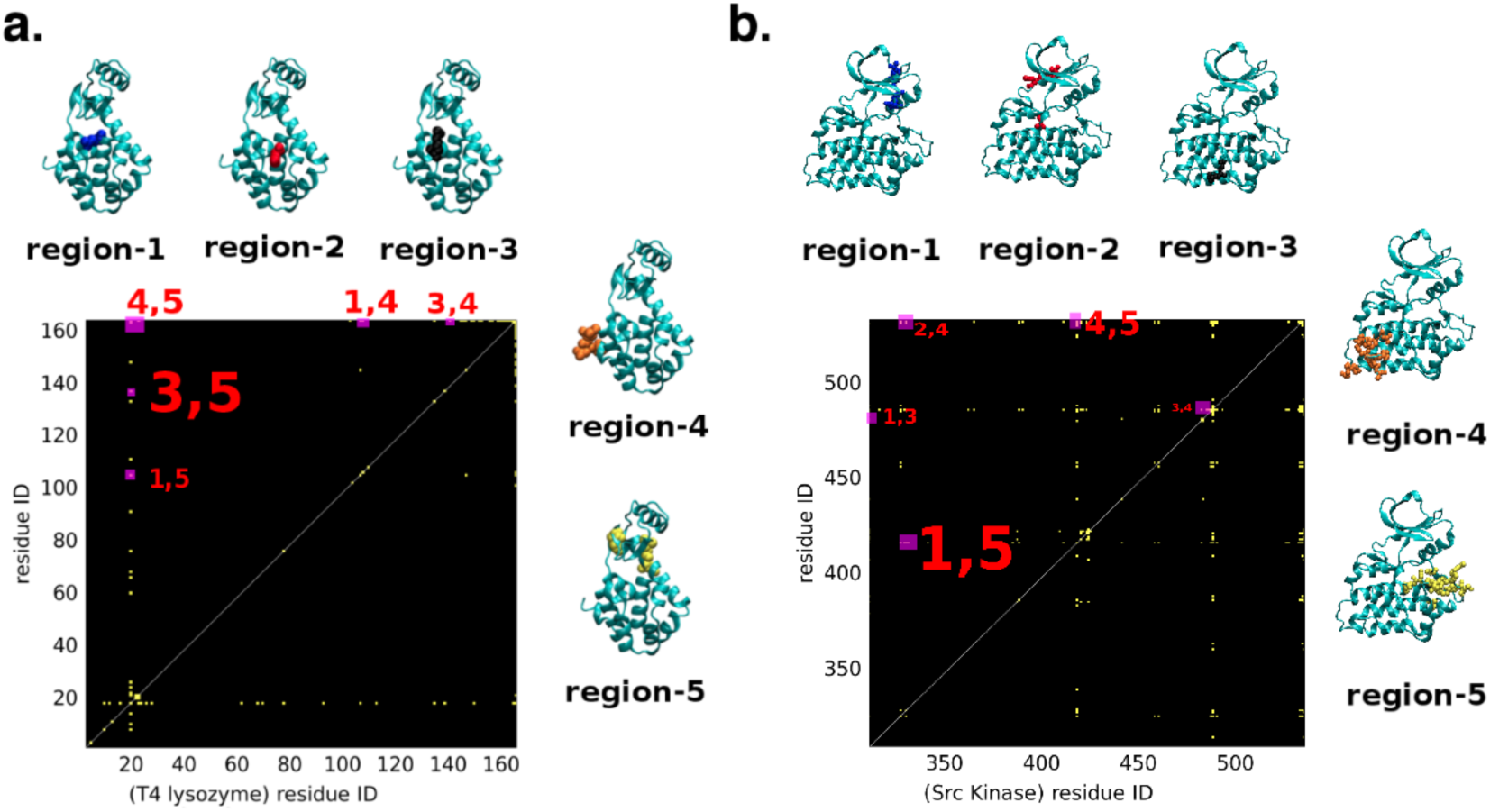
NAP correlation function plotted as a colour map spanning residue ids where correlation values larger than a threshold (= 4 percent of the maximum value) are shown in yellow while the rest is kept black; **a** T4Lys, **b** Src kinase. Highly correlated, spatially distant regions are sites for possible allosteric modification. Such pairs of sites are numbered in both cases with the font sizes corresponding to the relative value of the correlation (see also *SI Text*). The numbers correspond to the same regions as in Fig. 4. It is observed for T4Lys that the residue 18 from region 5 (yellow) has the highest correlation with the residue 137 in region 3 (black) as demonstrated by the largest font. In case of Src kinase, the residues 420 and 310, from regions 5(yellow) and 1(blue) respectively, have the highest correlation.

For Src kinase too, as shown in Fig. 5**b**, we could identify five regions of high NAP correlations at distant locations. For example, residue 416 (Tyr) of region **5** of src kinase has a very high NAP correlation with residue 328 (Val) (an element of Helix-*α*c) of region **1** (labelled **(1,5).** This is supported by a previous report^49^ of coupling between the activation loop and Helix-*α*c. A hallmark of active conformation of kinase^50^ is the autophosphorylation of the activation loop (404 - 432) at the residue 416 (Tyr) and”*α*C-in” conformation of Helix-*α*c. Analysis of non-affinity shows that the softest non-affine modes (video V4 in supporting info) at residue 416 and residue 328 in our simulations act as incipients for residue 416 (Tyr) protruding outward from the activation loop region, thereby becoming more exposed for the phosphorylation at this site (this site being tyrosine which is favourable for phosphorylation), and at the same time inward rotation of Helix-*α*c. In summary, our analysis based upon NAP-susceptibility and NAP spatial correlation function points towards allosteric contact between Helix-*α*c (helix undergoing inward rotation) and activation loop of kinase (phosphorylation site Tyr416 getting exposed).

## Multiple active sites in Cytochrome P450

The family of Cytochrome P450 proteins are quite versatile and are present in many different human tissues performing many tasks such as oxygen metabolism by binding to Heme, breakdown of toxins and hormones, synthesis of cholesterol^51^ etc. They are often associated with intracellular membranes such as in mitochondria and the endoplasmic reticulum. Extensive studies using sequence-based co-evolutionary analysis and anisotropic thermal diffusion MD simulations have identified four principal active regions in P450 related to membrane association, catalytic activity, Heme binding and dimerization.^52^

Our results for the NAP susceptibility is shown in Fig. 6. Remarkably, we can also identify the same regions as obtained in the earlier study,^52^ although the variant of P450 used by us is somewhat different. A complete analysis of all the different binding and allosteric processes of P450 analysed using the NAP values, eigenvectors and correlation functions is much beyond the scope of the present work and will be published elsewhere.

**Figure 6:**
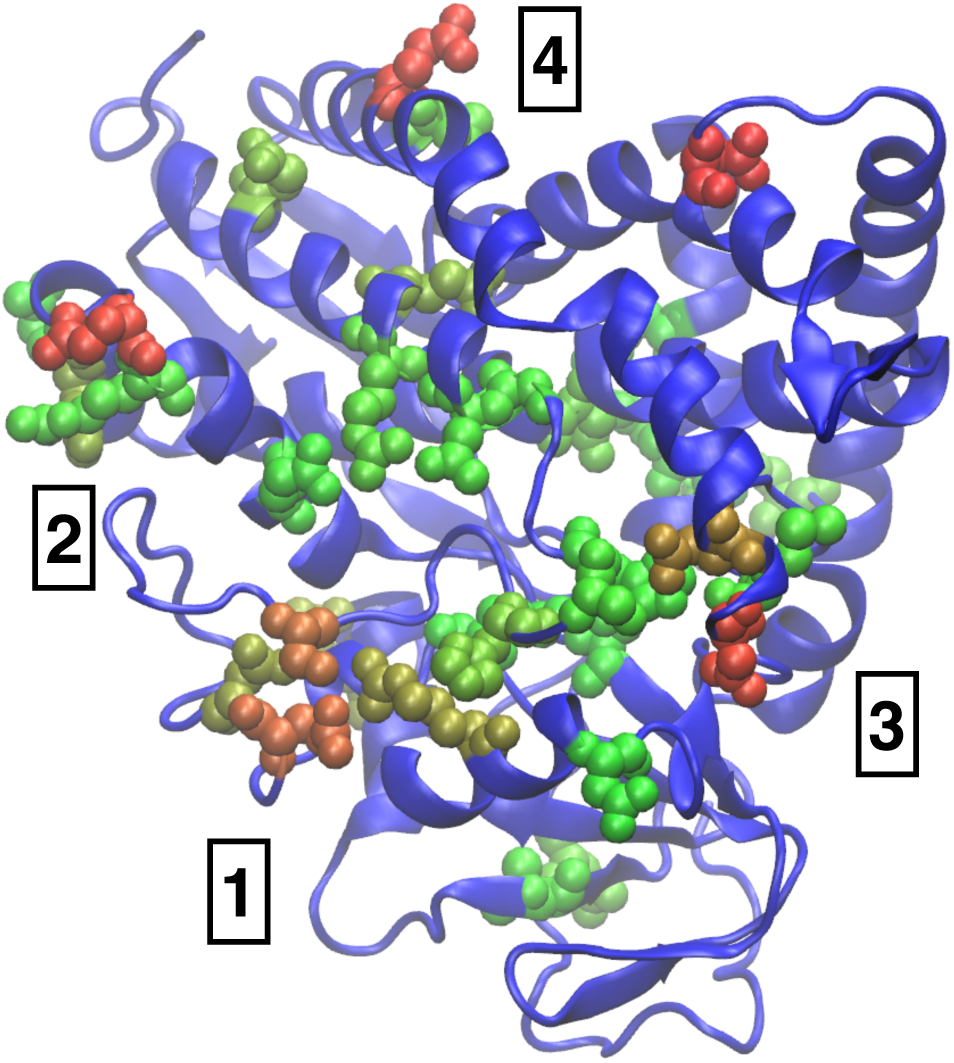
Relative NAP susceptibility plotted with the same color scale as in Fig 4 for Cytochrome P450. We have identified four regions of high activity. Region 1 and 2 are associated with membrane binding and catalytic activity respectively while 3 and 4 may correspond to Heme binding and dimerisation.

## Discussion and conclusion

Pinpointing sites of catalytic activity, binding of ligands or other functionally important conformational changes in proteins is a challenge when such displacements in protein residues are subtle and easily masked by background noise. Accurate “order parameters” or collective variables need to be designed to study such activity. It is often unclear how such collective variable may be defined. Drawing from ideas which were first demonstrated in crystalline solids^27–29^ we have defined a new collective variable, NAP, which quantifies non-affine thermally excited displacements of proteins. Firstly, we show that the susceptibility of different locations of a protein to non-affine displacements can be used as an indicator to determine regions which are important for binding events. Secondly, from the analysis of the NAP susceptibility values we conclude that the regions with higher values of NAP susceptibilities are also those which contain a dominant non-affine mode whose eigenvalue is separated from the those of the the rest of the modes by a large gap. The eigenvalue spectrum of the non-affine modes is also highly non-random. The eigenvector corresponding to this dominant mode involves atomic displacements that are the ones prone to act as binding gate-ways and the time spent by a ligand during a binding event at a particular residue is determined by the NAP susceptibility of this region. We also show that a positive correlation exists between the NAP value, its susceptibility and the magnitude of the gap in all the proteins studied by us.

Our analysis is fundamentally different from principal component analysis (PCA) ^53–55^ or other PCA based methods^56^ which are used extensively for analysing protein configurations obtained from simulations. The restriction to the coarse graining volumes Ω*_i_* makes the analysis local and the projection to the non-affine sub space guarantees that all trivial motions are filtered out. Finally, the large gap in the non-affine excitation spectrum between the dominant eigenmode and the rest ensures that there is very little mixing of the modes. As an example, recall the (un-)twisting motion of the helices observed in T4Lys as shown in the supplementary video V1 (see *SI Text*). Such a motion requires a special linear superposition of a large number of normal modes. The dominant PCA mode on the other hand cannot capture these motions (see supplementary FIg. S1 in *SI Text*) in detail although atomic displacements in the relevant regions are large. Unfortunately, these displacements are also contaminated by unimportant, affine motons which mask the gate opening feature.

The ability to compute NAP susceptibility by our approach helps us to unify two different proposed hypotheses of protein-ligand recognition, namely “conformational selection” ^57–62^ (in which protein’s intrinsic ability to shift to a ligand-preferred conformation guides the molecular recognition) and “induced fit” (in which ligand induces the protein to change its conformation). Our calculations show that these ideas are not in conflict but follows from standard fluctuation-response behaviour connecting local fluctuations of non-affine part of atomic displacements to the response of the protein to the conjugate local non-affine field.^28^ This is a simple consequence of a fluctuation response relation viz. 〈*χ*〉 = 〈*χ*〉_0_ + *h*_*χ*_〈(*χ* − 〈*χ*〉)^2^〉_0_ if we identify ligands with local fields *h*_*χ*_ which generate NAP in proportion to their susceptibilities at *h*_*χ*_ = 0. The ligand, in this case acts as such an “external” field and the response of the protein to the field is proportional to the fluctuations of local NAP in its apo state without the field (ligand).

Additionally, we see that the spatial correlation of NAP among various residues can be used to discover sites of possible allosteric control. NAP susceptibilities and correlations may be obtained readily from simulations of the apo protein without the ligand and come at a fraction of the cost needed for detailed simulations of ligand binding events. This implies that our proposed method is extremely well suited to be adapted for high throughput searches of possible protein ligand pairs and allosteric control.

It is to be noted that In our work we have focussed on the projection of the displacements to the non-affine subspace which is orthogonal to local strain.^46^ Since binding hotspots feature large displacements, both strain and strain fluctuations may be large together with large NAP values. NAP however, quantifies the error made in trying to fit the local displacements to an affine model and therefore large NAP values also signify that a description in terms of strain becomes invalid in those regions.

## Acknowledgement

We thank Madan Rao, Satyavani Vemparala and Pratyush Tiwari for discussions. HK is supported by Lundbeckfonden.

## Regions of Proteins involved in allostery

### L99A T4 Lysozyme, PBD ID 3MDV

The following are the 5 regions of T4Lys with high NAP susceptibiity:

| regions | residues | description |
| --- | --- | --- |
| **1** | 105 | part of helix-5 |
| **2** | 102 | part of helix-5 |
| **3** | 137 | part of helix-8 |
| **4** | 162 163 164 | — |
| **5** | 18 66 68 | — |

The relative values of the spatial NAP-correlation for the 5 locations shown in Fig.4(a) for T4Lys are as follows:

| regions | residues | Correlation function value |
| --- | --- | --- |
| **1,5** | 18,105 | 0.55 |
| **3,4** | 133,164 | 0.67 |
| **1,4** | 105,164 | 0.71 |
| **4,5** | 18,164 | 0.75 |
| **3,5** | 18,137 | 1.00 |

### Src kinase, PBD ID 1Y57

The following are the 5 regions of Src-kinase with large NAP susceptibility

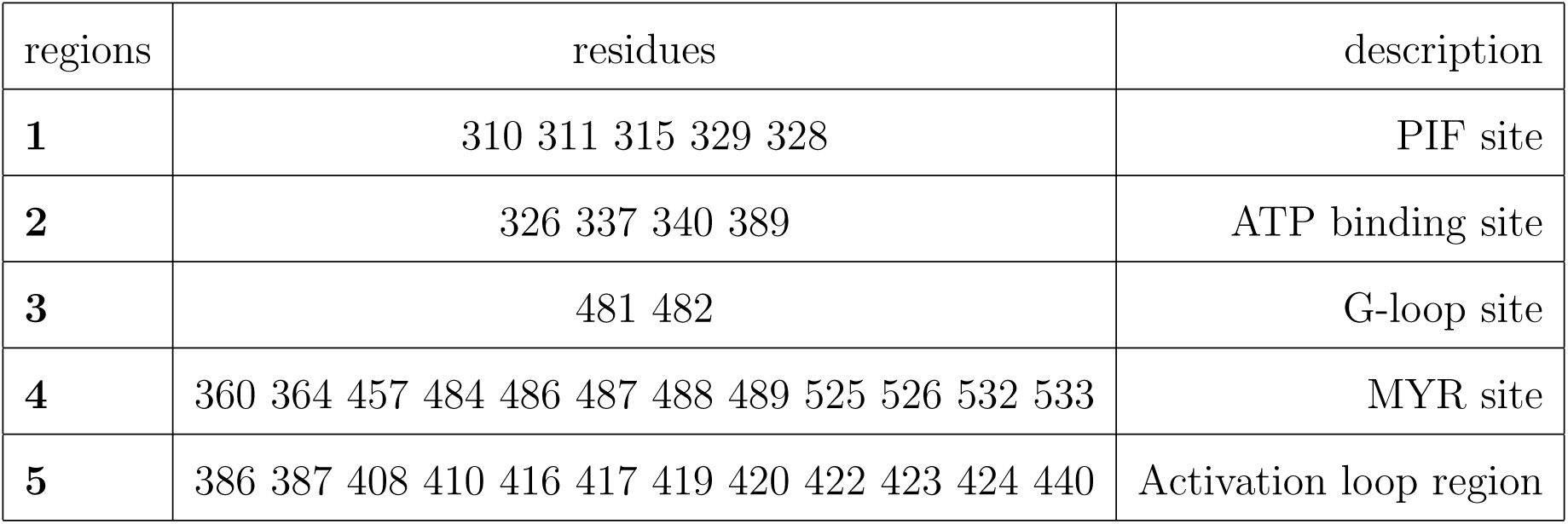

Src kinase, PBD ID 1Y57 The following are the 5 regions of Src-kinase with large NAP susceptibility
| regions | residues | description |
| --- | --- | --- |
| **1** | 310 311 315 329 328 | PIF site |
| **2** | 326 337 340 389 | ATP binding site |
| **3** | 481 482 | G-loop site |
| **4** | 360 364 457 484 486 487 488 489 525 526 532 533 | MYR site |
| **5** | 386 387 408 410 416 417 419 420 422 423 424 440 | Activation loop region |

The relative values of the spatial NAP-correlation for the 5 points shown in Fig.4(b) for Src-Kinase are as follows:

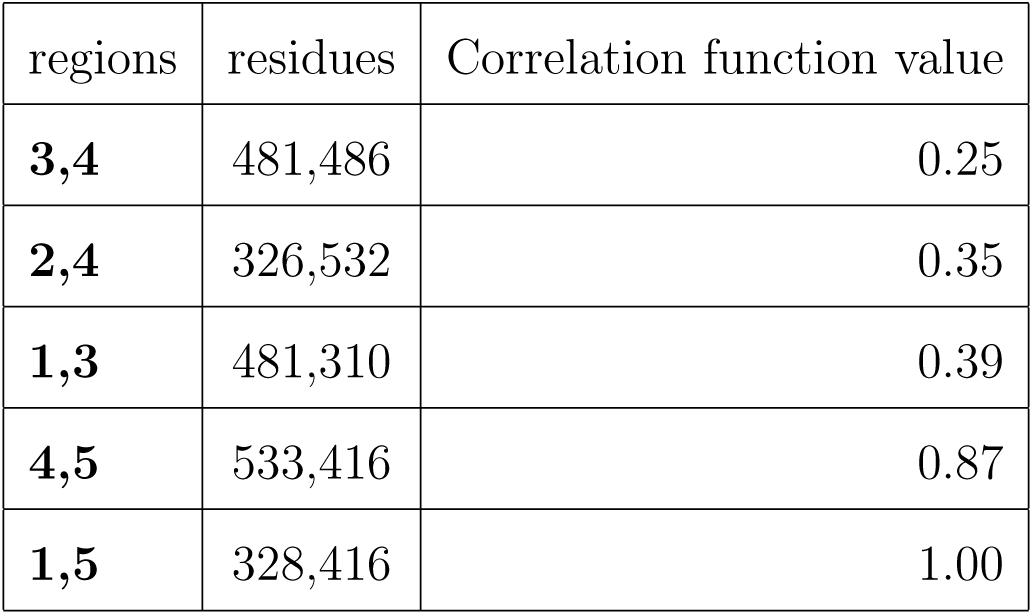

| regions | residues | Correlation function value |
| --- | --- | --- |
| **3,4** | 481,486 | 0.25 |
| **2,4** | 326,532 | 0.35 |
| **1,3** | 481,310 | 0.39 |
| **4,5** | 533,416 | 0.87 |
| **1,5** | 328,416 | 1.00 |

## Description of Videos

We have included four videos which show prominent non-affine displacements. In each of these videos we have applied the eigenvectors corresponding to the largest eigenvalue of the local PCP matrix to cause displacements of the atoms corresponding to the native structure of the protein. Note that these are not MD simulations.

1. The video V1 demonstrates the opening of the helices 4 and 6 in order to enable the ligand to enter the binding site using the pathway depicted in Fig. 2 **a** of the manuscript. Note the twisting of the helices in this video and in V2 below.
2. The video V2 demonstrates the opening of the helices 7 and 9 in order to enable the ligand to enter the binding site using the pathway depicted in Fig. 2 **b** of the manuscript.
3. The video V3 demonstrates the non-affine modes for the residues having high spatial NAP correlation shown in Fig.4(a), thereby demonstrating the allosteric contact between regions 3 and 5 of T4Lys. The color scheme used is same as Fig.4(a). The loop in vicinity of region 3 of which ARG137 is a part is shown in black. and the loop of which TYR18 is a part, being a neighbourhood of region 5, is shown in yellow. The Alfa-carbon atoms of the two residues (TYR18, ARG137 of regions 5 and 3 respectively) having high spatial NAP correlation are shown in pink solid bubbles. Here we see the residue 22(GLU) yellow, initially forming a hydrogen bond with residue ARG(137) black moving apart from residue 137 black. Thus, the non-affine modes at residue 18 and 137 act as nucleating precursors which tend to take T4Lys molecule from the closed conformation to open conformation.
4. The video V4 demonstrates the non-affine modes for the residues having high correlation shown in Fig.4(b), thereby demonstrating the allosteric contact between regions 1 and 5 of SRC-Kinase. The color scheme used is same as Fig.4(b). The activation loop alongwith residue Tyr416 being a part of region 5 is shown in yellow. and the Alfa-c helix alongwith residue 328 being a neighbourhood of region 1 is shown in blue. The Alfa-carbon atoms of the two residues (Tyr416, Val328 of regions 5 and 1 respectively) having high spatial NAP correlation are shown in pink solid bubbles. Here we see the residue 416 protruding outside from the activation loop region thereby becoming more exposed and facilitating the phosphorylation. At the same time, the Alfa-c helix is seen to be undergoing an inward rotation. Non-affine modes analysis show that the incipients for the outwards protruding of Tyr416 and incipients for the inwards rotation of Alfa-c helix have an allosteric contact.

| NAME | Phenomenon |
| --- | --- |
| **V1** | opening-up of helices 4 and 6 |
| **V2** | opening-up of helices 7 and 9 |
| **V3** | close to open configuration |
| **V4** | allosteric contact: activation loop and Helix-αc |

## Relation with PCA

In our work non-affine modes of fluctuations are projections from the total displacements **Δ** by allowing the operator or matrix P to act on **Δ** i.e. we calculate P**Δ.** We therefore perform a normal mode analysis over this projected subspace containing solely the non-affine modes of deformation by calculation of the local, projected Sample Correlation Matrix (SCM) 〈(P**Δ**)(P**Δ**)*^T^*〉 where angular brackets represent ensemble average for the trajectories of atoms in the ligand free protein. Carrying out this ensemble average we obtain the correlator as PCP. Here C = 〈**ΔΔ***^T^*〉 is an SCM for ordinary (total) displacement **Δ** undergone by the atoms within the coarse-grain volume.

We have shown that from a normal mode analysis of P**Δ** we obtain the eigenvalues and eigenvectors describing the most important and relevant dynamical modes for protein function. The softest eigenmodes or the eigenvectors corresponding to the largest eigenvalues are demonstrated in the videos provided in the supporting material (videos V1-V4). The modes of fluctuations shown in these videos indeed demonstrate that these non-affine modes are actually responsible for bringing about the important functions like the opening-up of the gateways leading to the binding pockets and the allosteric interactions between the distal functional sites.

This projection to the non-affine subspace is crucial. An ordinary principal component analysis (PCA) keeps the full displacement correlator of the whole protein without the projection. In Fig. S1 we show the result of this calculation for T4Lys. Understandably, none of the relevant conformation changes related to gate opening can be seen. The displacements are overwhelmingly affine.

**Figure S1:**
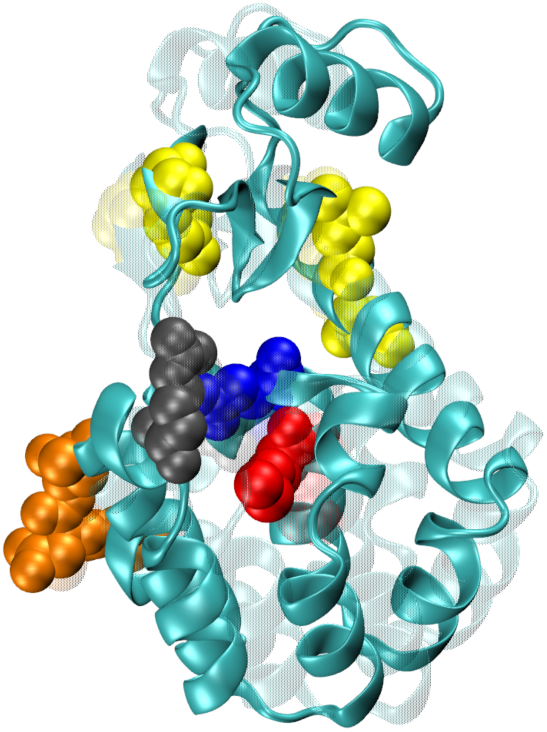
The solid cyan image is the original crystal structure of T4 Lysozyme. The ghost cyan image is the final orientation of the protein after the application of the most prominent (highest eigenvalue) eigenmode obtained from the Principal Component Analysis. For comparison with our NAP susceptibility calculations, the regions of large susceptibility are highlighted with the same color scheme as in Fig.5 of the manuscript. Note that now no helix twisting motion can be seen in region 1 and 2. The PCA displacements are seen to be mostly affine.

